# Identification of Putative Cell Wall Synthesis Genes in *Betula pendula*

**DOI:** 10.1101/2020.05.21.107581

**Authors:** Song Chen, Xiyang Zhao, Su Chen

**Author notes:** **Correspondence:** (Su Chen).

## Abstract

Cellulose is an essential structural component of the plant cell wall and is an important resource for the production of paper, textiles, bioplastics and other biomaterials. The synthesis of cellulose is among the most important but poorly understood biochemical processes, which is precisely regulated by internal and external cues. Here we identified 46 gene models in 7 gene families which encoding cellulose synthase and related enzymes of *Betula pendula*, and the transcript abundance of these genes in xylem, root, leaf and flower tissues also be determined. Based on these RNA-seq data, we have identified 8 genes that most likely participate in cell wall synthesis. In parallel, a gene co-expression network was also constructed based on transcriptome sequencing.

**Funding:** This work was supported by the National Natural Science Foundation of China, grant number 31870659, The Fundamental Research Funds for the Central Universities, grant number 2572019CG08 funded this research and Heilongjiang Touyan Innovation Team Program (Tree Genetics and Breeding Innovation Team).

**Conflicts of interest / Competing interests:** The authors declare that the research was conducted in the absence of any commercial or financial relationships that could be construed as a potential conflict of interest.

**Consent to participate:** Not applicable.

**Consent for publication:** Not applicable.

**Availability of data and material:** All data generated or analyzed during this study are included in this published article.

**Code availability:** Not applicable.

**Authors’ contributions:** All authors contributed to the study conception and design. Material preparation, data collection and analysis were performed by Song Chen. Conceived and supervised were performed by Xiyang Zhao and Su Chen.

## 1 Introduction

Silver birch (*Betula pendula*) is a medium-sized deciduous tree that owes its common name to the white peeling bark on the trunk. This species is native to Europe and parts of Asia, and the range extends into Siberia, China, and southwest Asia in the mountains of northern Turkey, the Caucasus, and northern Iran (Hynynen et al. 2010). Recently, the pace of genome sequence generation has increased and the assembled sequences of *B. pendula* have become publicly available, which can help us understand this species at the genome expression level (Salojärvi et al. 2017).

The cell periphery of higher plants is usually surrounded by the cell wall. Plant cell walls are complex networks of polymers that provide protection and structural properties to the cells (Buchanan et al. 2015). The cell wall mainly includes four major chemical polymers: cellulose, hemicellulose, lignin and pectin. The most characteristic of the xylem cell wall is higher cellulose content, cellulose contains a large amount of cellulose synthase, which visualized as symmetrical rosettes (Kimura et al. 1999). The only known component of cellulose synthase in plants is the CESA protein, which was originally isolated and identified in cotton (Pear et al. 1996). Subsequent analysis of the *A. thaliana* genome revealed that a total of 10 genes encode CESA proteins with 64% average sequence identity (Holland et al. 2000; Richmond 2000). *Populus trichocarpa* has 18 CESA genes (Djerbi et al. 2005), *Hordeum vulgare* has 8 (Burton et al. 2004) and *Zea mays* has 12 (Appenzeller et al. 2004).

As an important tree species in papermaking, understanding the cellulose synthesis pathway of *B. pendula* will greatly contribute to its use in industrial production. In this study, we identified the genes that likely encode cellulose synthase and related enzymes during cell wall synthesis in *B. pendula*, which will serve as a basis for further gene functional studies.

## 2 Materials and methods

### 2.1 Identification of *B. pendula* cell wall synthesis genes

The *B. pendula* genome (Salojärvi et al. 2017) and genomic structure information (GFF) were downloaded from the CoGe comparative genomics platform. The putative cellulose synthase genes were first identified by BLASTP v2.9.0 (Camacho et al. 2009) with the *A. thaliana* cellulose synthase genes as queries (E-value ≤ 1E-5). We then further manually examined these putative cell wall synthesis genes using the Conserved Domain Database of NCBI (Marchler-Bauer and Bryant 2004) to confirm if they were correctly annotated, and divided them into seven subgroups based on their functional type in *A. thaliana*. Molecular weight and theoretical isoelectric point were analyzed by the ExPaSy Compute pI/Mw tool (Gasteiger et al. 2003), and the chromosomal location of the *B. pendula* cell wall synthesis genes was visualized by using TBtools v0.67 (Chen et al. 2018).

### 2.2 Phylogenetic analyses of *B. pendula* cell wall synthesis genes

To investigate the phylogenetic relationships of the cellulose synthases (CESAs) and cellulose synthase-like proteins (CSLs), the phylogenetic tree was constructed for every subgroup. The multiple sequence comparison was performed by MUSCLE v3.8.1551 (Edgar 2004) with default parameters, and the constraint maximum likelihood phylogenetic trees of each subgroup were then be generated by RAxML v8.2.12 (Stamatakis 2014) with 1,000 bootstrap trials. The model was selected for the GAMMA model and visualized by iTOL v5 (Letunic and Bork 2019).

### 2.3 RNA-seq expression analysis of *B. pendula* cell wall synthesis genes

We downloaded the transcriptome data (PRJNA535361) (Chen et al. 2019) from the NCBI SRA database to investigate the expressional patterns of *B. pendula* cellulose synthase genes in different tissues. The clean reads of three replicates per tissues were aligned to the *B. pendula* transcriptome by using STAR v2.7.3a (Dobin et al. 2013), and the accurate transcript quantification was estimated by using RSEM v1.3.3 (RNA-seq by Expectation-Maximization) pipeline (Li and Dewey 2011) with paired-end sequencing mode. The normalized expression value was all selected as TMM (trimmed mean of M-values).

### 2.4 Transcription factor regulatory networks in *B. pendula* cell wall synthesis

The transcription factors of *B. pendula* were identified by PlantTFcat (Dai et al. 2013), and the NCBI CD-search (Marchler-Bauer et al. 2015) to be used determine whether they were correctly annotated. To perform the weighted correlation network analysis (WGCNA) between cell wall synthesis genes and transcription factors, we used the WGCNA R package v1.69 (Langfelder and Horvath 2008) to construct the co-expression network. The TMM value from different tissues of *B. pendula* was as input expression data for this software, and only genes with TMM values larger than 10 for all samples were kept. The threshold power (β) value was determined to be 13 from pickSoftThreshold output, and the Pearson algorithm is then be used to calculate the correlation coefficient. Finally, the co-expression network was generated block wise using the WGCNA function blockwiseModules with the following settings: TOM-type, unsigned; mergeCutHeight, 0.15; deepSplit, 2; minModuleSize, 30; and eventually visualized by the Cytoscape v3.8.0 (http://cytoscape.org/).

## 3 Results

### 3.1 Identification of *Betula pendula* cellulose cell wall synthesis genes

A total of 29,439 coding genes in *B. pendula* genome (Salojärvi et al. 2017) were used to identify putative cell wall synthesis genes. In total, 46 gene models (Table 1) in 7 families were identified as putative cell wall synthesis genes in *B. pendula* genome. The 46 genes encode 10 cellulose synthase proteins (CESAs) and 36 cellulose synthase-like proteins (CSLAs, CSLBs, CSLCs, CSLDs, CSLEs and CSLGs) in 7 families. Among these families, *CESA* was the predominant cellulose synthase gene family and contains seven members. The rest of the gene families all belong to the cellulose synthase-like family, *CSLG* was the largest cellulose synthase-like family containing eleven members, while *CSLA* was the smallest family with only three members. We then applied quantitative criteria to assign the genes likely to be cell wall synthesis genes based on transcript abundance and specificity. The tissue-specific expressional data include xylem, roots, leaves and flowers, and we calculated the expression of the 46 identified genes. A total of 8 genes showed that expression in the xylem was higher than the expression in both flower and leaf. These genes were identified as the cell wall synthesis genes *BpCESA4*, *BpCESA9*, *BpCESA10*, *BpCSLA2*, *BpCSLA3*, *BpCSLC1*, *BpCSLC4* and *BpCSLD4*.

**Table 1.**
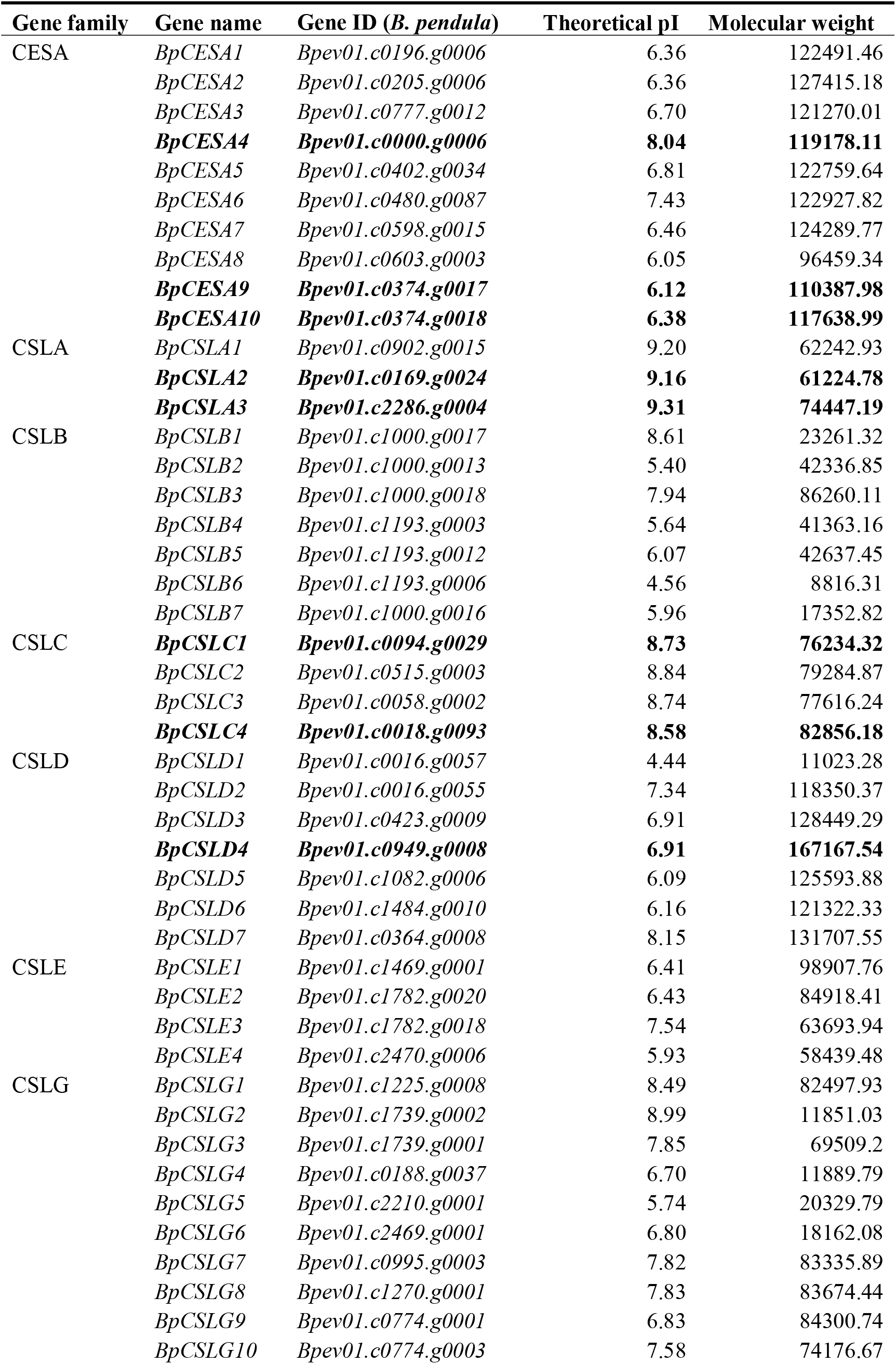
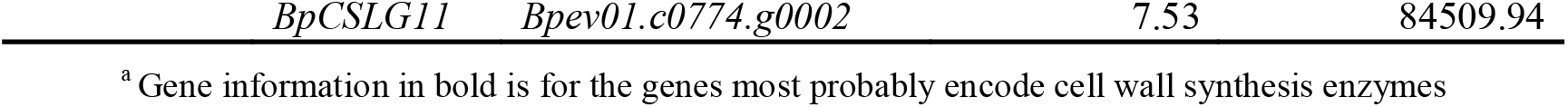
Putative *B. pendula* cellulose synthase genes in 7 gene families

### 3.2 Chromosomal location and gene duplication

Cell wall synthesis was mainly composed of cellulose synthases (CESAs) and cellulose synthase-like proteins (CSLs), so we investigated the formation of *CESAs* and *CSLs* based on the chromosomal location and intra-genome syntenic information. Similar to the *A. thaliana*, the multiple *BpCESAs* were scattered across the *B. pendula* genome and mapped in 13 of the 14 chromosomes (Figure 1). The *BpCESAs* were concentrated on Bpe_Chr6, Bpe_Chr7, Bpe_Chr8, Bpe_Chr9, Bpe_Chr10 and Bpe_Chr11, with one or two genes per chromosome. The *BpCSLs* were scattered on 13 chromosomes except for Bpe_Chr5, and we found that some *BpCSLs* were organized into duplicated blocks, such as *BpCSLB1-7* on Bpe_Chr2, *BpCSLG2-7* on Bpe_Chr14 and *BpCSLG8-10* on Bpe_Chr1. This situation always originated from the duplicative transposition.

**Figure 1.**
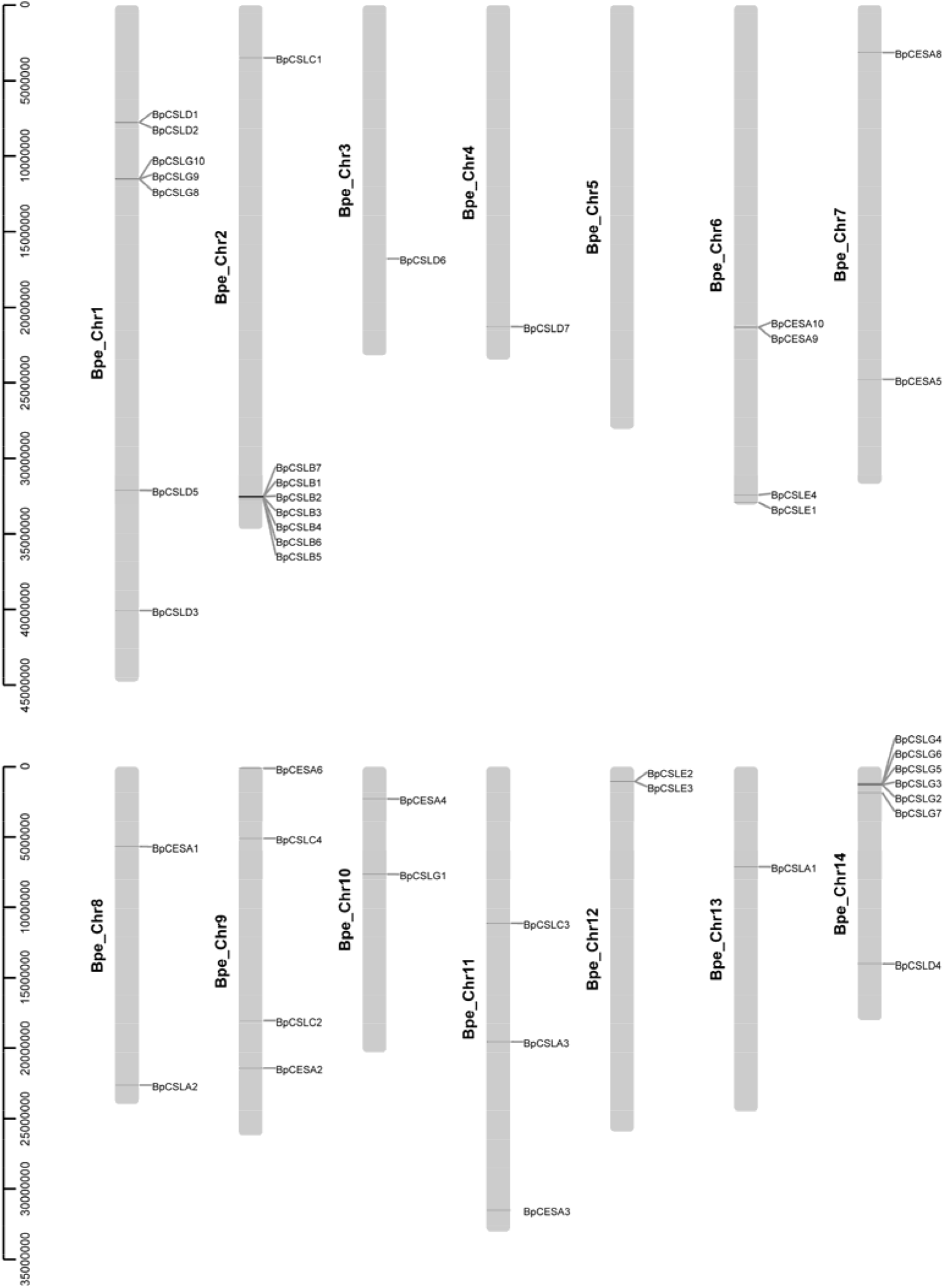
The chromosomal location of *B. pendula* cell wall synthesis genes. The silver line represents the chromosome of *B. pendula*, and the black line represents the relative location of CESA and CSL genes on the chromosome.

### 3.3 Cellulose synthase (CESA) gene family

Cellulose was the principal ingredient of the cell walls in *B. pendula*, small microfibrils were crystallized by 36 tails of H-bonded-β-1,4-Glc chains catalyzed by cellulose synthases (Joshi 2003). Thus, cellulose synthase was one of the indispensable glycosyltransferases in plants, which plays a crucial role in regulating cell wall cellulose synthesis and plant cell morphogenesis.

We identified 11 *BpCESAs* in the *B. pendula* genome, of which *BpCESA4*, *BpCESA9* and *BpCESA10* were abundant in xylem (Figure 2). *BpCESA4* was the highest expressed gene in the root and xylem of the CESA family. The most similar protein to BpCESA4 was AtCESA4 in *Arabidopsis thaliana*, which confers plant resistance to bacterial and fungal pathogens while encoding a cellulose synthase. Handakumbura et al. (2013) reported that AtCESA4 loss-of-function mutants of *A. thaliana* and *Oryza sativa* have weak stems and thin or irregular cell walls. The protein most similar to BpCESA9 and BpCESA10 was AtCESA8 in *A. thaliana*, Glass et al. (2015) reported that endo◻1,4◻glucanases AtGH9B5 and AtGH9C2 can impact cellulose β crystallization and plant cell wall development by influencing cellulose synthase AtCESA8. In addition, Kim et al. (2014a) reported that transcription factor AtMYB46 can directly regulate the secondary wall-associated cellulose synthase AtCESA4 and AtCESA8 in *A. thaliana*.

**Figure 2.**
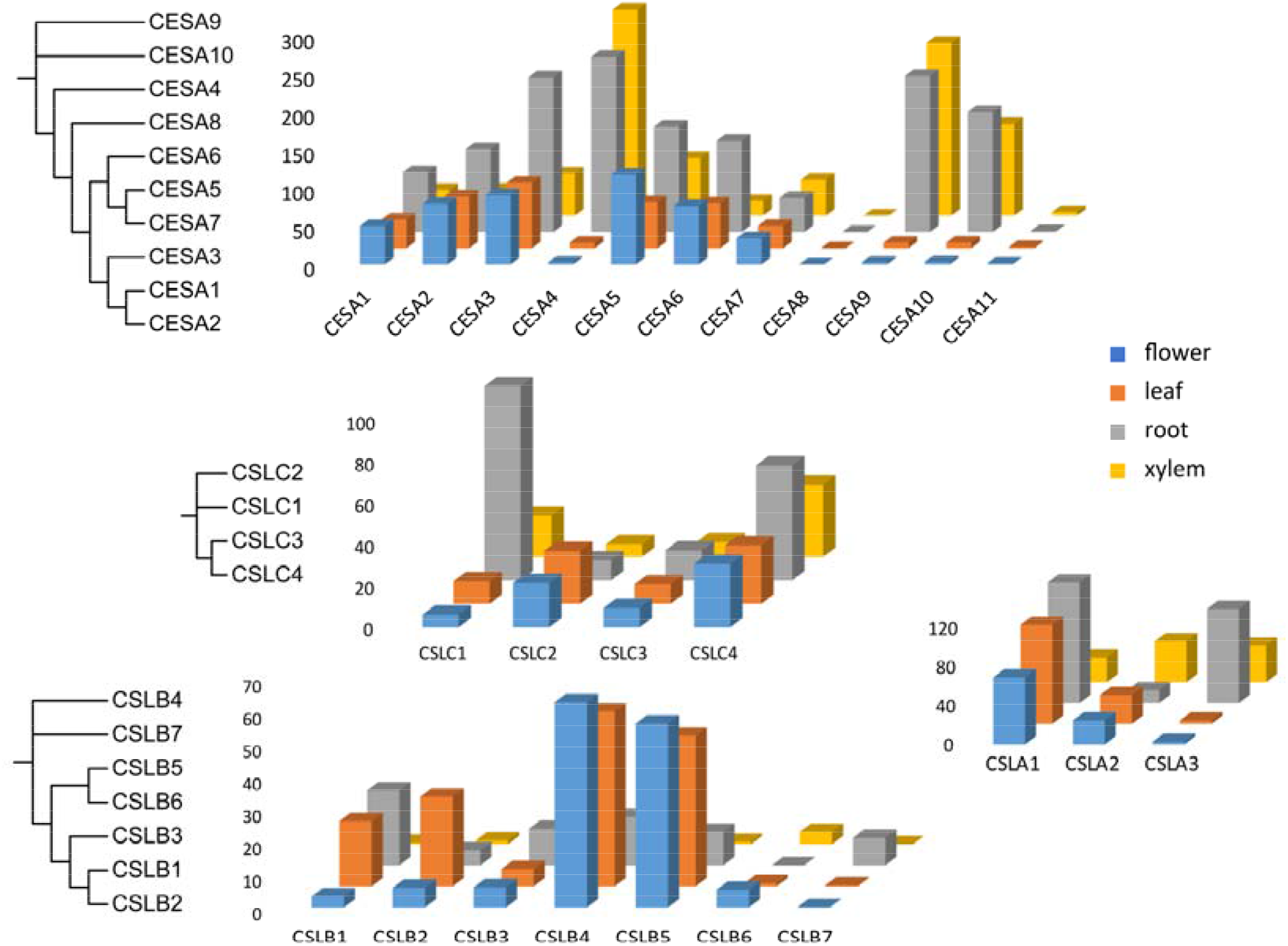
Tissue-specific expression profiles and phylogenetic analysis of *CESA*, *CSLA*, *CSLB* and *CSLC* families in *B. pendula*. The expression was analyzed in three independent biological replicates of each tissue, and the phylogenetic tree (1,000 bootstraps) was constructed by RAxML using the maximum likelihood algorithm.

### 3.4 Cellulose synthase-like (CSL) gene family

The cellulose synthase-like gene family was divided into six families, which were CSLA, CSLB, CSLC, CSLD, CSLE and CSLG. The function of the CSL family was still explored now and only a few reports were available. Jensen et al. (2012) reported that the CSL genes is associated with hemicellulose synthesis, Schreiber et al. (2014) and Doblin et al. (2009) reported that cellulose synthase-like protein CSLFs and CSLHs mediate the synthesis of cell wall (1,3)(1,4)-β-D-Glucans, but the vast majority of CSL genes functions require further study.

We identified 38 *BpCSLs* in the *B. pendula* genome of which 5 genes were abundant in xylem (Figures 2 and 3). They were *BpCSLA2*, *BpCSLA3*, *BpCSLC1*, *BpCSLC4* and *BpCSLD4*, respectively. BpCSLA2 and BpCSLA3 were most similar to AtCSLA9 in *A. thaliana*. Expression of *CSLs* in *A. thaliana* cells revealed that AtCSLA glycosyltransferases can encode cell wall glucomannan and intervention the progression of embryogenesis (Goubet et al. 2009; Liepman et al. 2005). In addition, Kim et al. (2014b) reported that transcription factors AtNAC41, AtbZIP1 and AtMYB46 can directly regulate the expression of *AtCSLA9* in *A. thaliana*. The most similar protein to BpCSLC1 was AtCSLC4 in *A. thaliana*, which encodes a protein similar to cellulose synthase and its mRNA can mobile in cell-to-cell. The 1,4-beta-glucan synthase AtCSLC4 can form the xylosylated glucan backbone with three xylosyltransferases AtXXT1, AtXXT2 and AtXXT5 in *A. thaliana* (Chou et al. 2012). Intriguingly, glucan synthase AtCSLC4 have opposite orientations in the Golgi membrane (Davis et al. 2010) with mannan synthase AtCSLA9, which may cause the functional differences between them. The most similar protein to BpCSLD4 was AtCSLD3 in *A. thaliana*, which part in the cell-wall synthesis of tip-growing root-hair cells (Park et al. 2011). Galway et al. (2011) reported that root hair-specific disruption of cellulose and xyloglucan in AtCSLD3 mutants.

**Figure 3.**
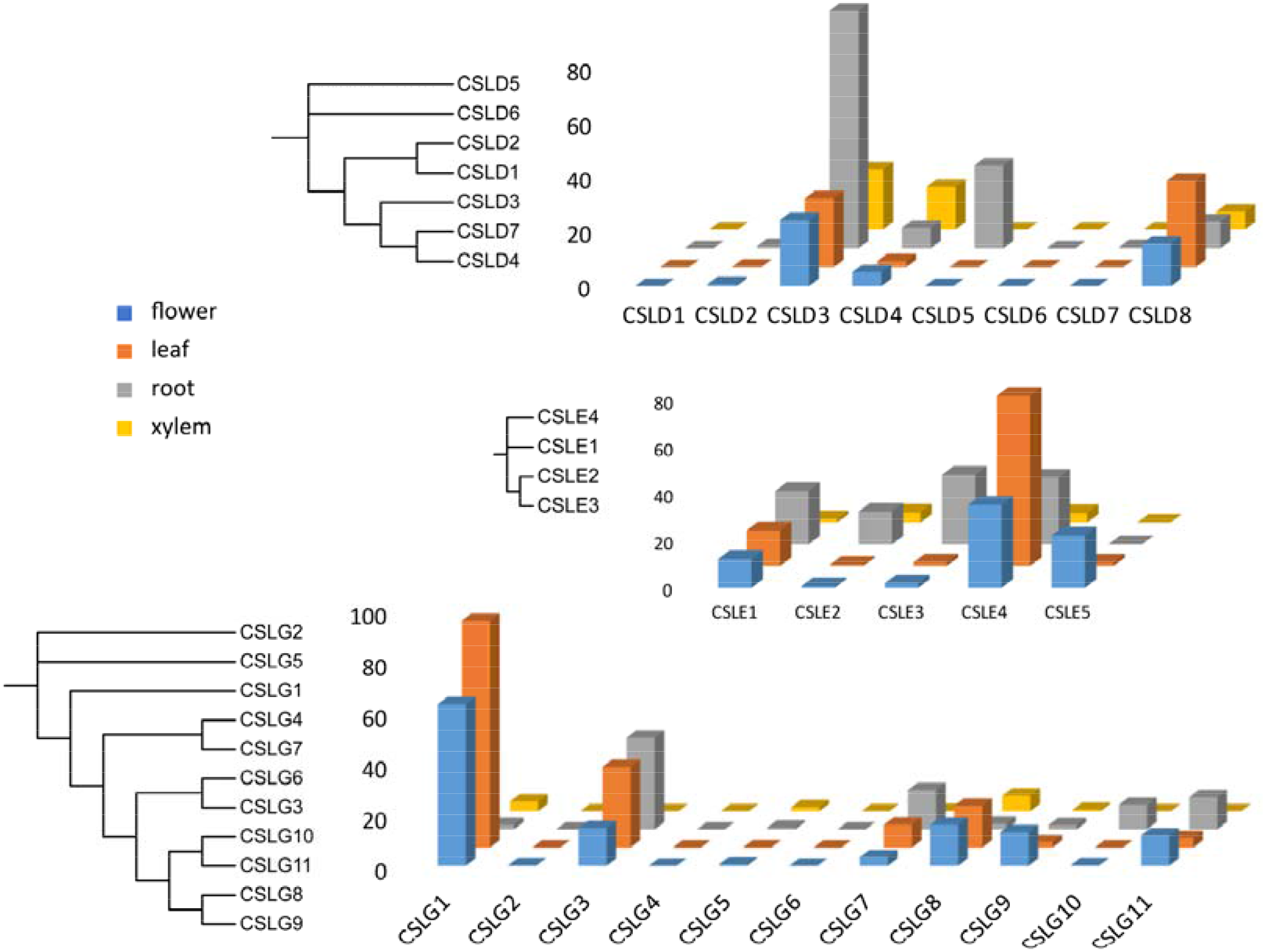
Tissue-specific expression profiles and phylogenetic analysis of *CSLD*, *CSLE* and *CSLG* families in *B. pendula*. The expression was analyzed in three independent biological replicates of each tissue, and the phylogenetic tree (1,000 bootstraps) was constructed by RAxML using the maximum likelihood algorithm.

### 3.5 Involvement of transcription factors in cell wall synthesis

Based on transcriptome sequencing data, we performed an extensive analysis between putative cell wall synthesis proteins and 2,816 transcription factors (Table S1) of *B. pendula*. The results showed that a total of 51 transcription factors were co-expressed with 6 cell wall synthesis proteins, which were BpCESA4, BpCESA9, BpCSLA2, BpCSLC1, BpCSLC4 and BpCSLD4 (Figure 4).

**Figure 4.**
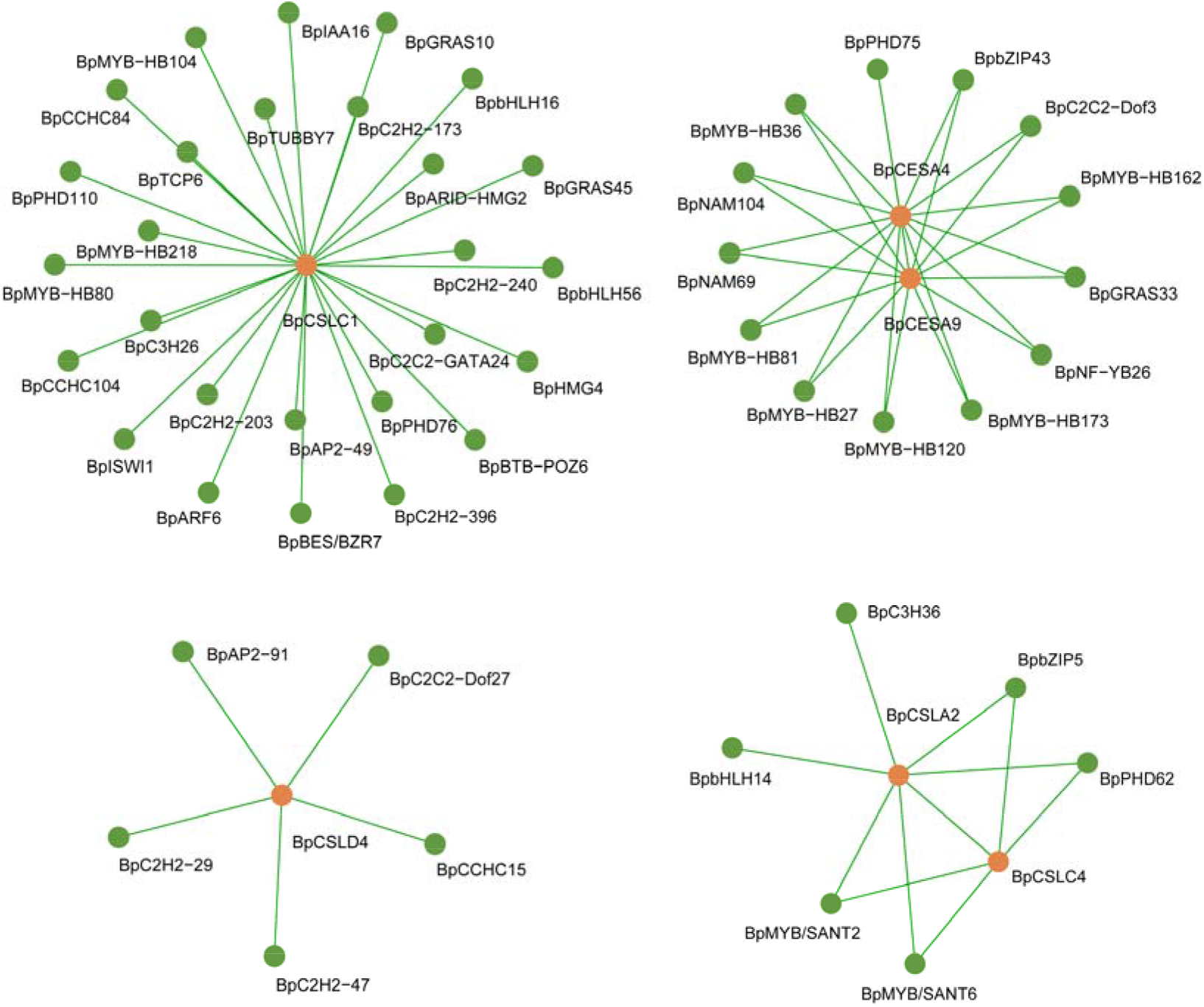
The transcription factor regulatory network calculated by WGCNA. The green dots were transcription factors, and the orange dots were cell wall synthesis genes.

The highest number of transcription factors were co-expressed with BpCSLC1, up to 27, including ARF, IAA and several other auxin-related transcription factors. BpARF6 was most similar to AtARF17 in *A. thaliana*, Yang et al. (2013) reported that AtARF17 is essential for primexine formation and pollen wall development. BpIAA16 was most similar to AtIAA16, which has transcriptional wiring with cell wall-related genes in *A. thaliana*, too (Mutwil et al. 2009). In addition to BpCSLC1, there was a co-expression relationship between BpCESA4 and BpCESA9, with 13 transcription factors regulating these two cellulose synthase genes. BpNAM69 was most similar to AtNAC43 (NST1) in *A. thaliana*, which is known to be involved in cellulose synthesis. Zhong et al. (2007) reported that inhibition of the expression of both *AtSND1* and *AtNST1* by RNA interference (RNAi) results in loss of secondary wall formation in stem fibers, and several fiber-associated transcription factor genes will be down-regulation in *A. thaliana*. BpMYB-HB162 was most similar to AtMYB83 in *A. thaliana*, Ko et al. (2014) reported that the AtMYB46/AtMYB83-mediated transcriptional regulatory program is a gatekeeper of secondary wall synthesis.

## 4 Discussion

In this study, we identified a total of 8 genes that most likely involved in cell wall synthesis in *B. pendula*, which should help elucidate the molecular mechanism of cellulose synthesis in *B. pendula*. These genes showed striking consistency compared to the cell wall synthesis genes in *P. trichocarpa*, demonstrating that the cellulose synthesis family is conserved during species evolution.

Cellulose synthesis requires the plant hormones, nitric oxide and cellulose synthase, and this coordinated control involves a multifaceted and multilayered transcriptional regulatory program. We can effectively modulation this process by promotes several enzymes expression, thereby changing the cellulose content of *B. pendula*. Oomen et al. (2004) reported that reducing of the cellulose content of *Solanum tuberosum* tuber by antisense expression of *StCESA3* clones. Zhong et al. (2003) reported that the *AtCesA7* mutant of *A. thaliana* has lower fiber cell wall thickness and cellulose content. However, the process of increasing cellulose content is not as simple as reducing it. Tan et al. (2015) reported that overexpressing *HvCESA* showed no increase in cellulose content or stem strength in *Hordeum vulgare*, despite the use of a powerful constitutive promoter. Previous studies (Doblin et al. 2002) have shown that individual CESA and CSL proteins play different roles in the synthase complex and require tightly regulated, so we need more complex strategies in the plant engineering of increasing cellulose content.

## 5 Conclusion

This study aims to provide information on *B. pendula* cell wall synthesis genes regarding their potential physiological roles and the molecular mechanism associated. In this study, we identified a total of 8 cell wall synthesis genes in *B. pendula*, which include 3 cellulose synthase genes and 5 cellulose synthase-like genes. And a gene co-expression network was constructed based on synthesis-related genes expression value. These analyses will help decipher the genetic information of the cell wall synthesis genes, elucidate the molecular mechanism of cellulose synthesis and understand the cell wall structure in *B. pendula*.

## 6 Data Archiving Statement

The RNA datasets used in the current study are available in the NCBI SRA (Sequence Read Archive) database (Accession No. PRJNA535361). The leaves, roots, xylem, and flowers of the two-year-old *B. pendula* were sampled and sequenced by Illumina HiSeq 2500.

## Supporting information

Table S1 Transcription factors of Betula pendula

